# The 11β-hydroxysteroid dehydrogenase type 1 inhibitor SPI-62 prevents morbidity in a mouse model of Cushing’s syndrome

**DOI:** 10.1101/2024.05.27.596076

**Authors:** Xingyu Pan, Qiongqiong Hou, Jing Su, Jiahui Xu, Min Li, Jiawen Li, Xuerou Shi, Caleb Schmicker, Tobias Magers, William M Bracken, David A. Katz

## Abstract

11β-hydroxysteroid dehydrogenase type 1 (HSD-1) inhibitors represent a potential therapeutic approach for patients with Cushing’s syndrome, autonomous cortisol secretion (ACS), or iatrogenic glucocorticoid (GC) excess. We assessed whether the HSD-1 inhibitor SPI-62 prevents GC-associated morbidity in a mouse Cushing’s syndrome model. Corticosterone (CORT) was administered to mice for 5 weeks. Animals who received CORT were randomized between vehicle and 3 SPI-62 regimens. A control group received neither CORT nor SPI-62. Body weight was measured daily to enable weight-based SPI-62 administration. Food consumption by pair-housed animals was reported weekly. Insulin sensitivity was evaluated by fasting homeostatic model assessment of insulin resistance. Adiposity and skeletal myopathy were measured using magnetic resonance imaging and post-mortem weights of fat depots and skeletal muscles. Grip strength was measured with a digital meter. Ambulation behavior in an open field maze was assessed. Post-mortem dermal thickness was quantified. Skin structure was evaluated by histology. CORT was associated with decreased insulin sensitivity, increased adiposity, skeletal myoatrophy, reduced grip strength, decreased dermal thickness, and skin structural changes. A trend of increased food consumption and an unexpected biphasic weight change were observed with CORT. SPI-62 attenuated all observed adverse effects in a dose-dependent manner. In general, results for the high SPI-62 regimen were similar as those in animals who received no CORT, suggesting that full HSD-1 inhibition should be maintained throughout a dose interval to mitigate the effects of glucocorticoid excess.

## Introduction

Glucocorticoid (GC) excess – whether of endogenous cortisol or a medicine such as prednisolone – is associated with considerable metabolic, muscular, dermal, and other morbidity. The canonical presentation of GC excess in patients with ACTH-dependent hypercortisolism, e.g., Cushing’s disease,includes centripetal obesity, plethoric moon facies, insulin resistance/diabetes, hypertension, hyperlipidemia, proximal myoatrophy and weakness, dermal thinning with striae and easy bruising, osteopenia, infection susceptibility, and mood, sleep, and cognitive disorders, as well as increased mortality (1–2). That morbidity and excess mortality risk is shared by patients with ACTH-independent hypercortisolism, e.g., autonomous cortisol secretion (ACS) associated with a benign adrenal adenoma which might affect as much as 1% of the population (3), as well as by millions of patients who rely on long-term use of GC medicines to control autoimmune diseases and other conditions. Those medicines account for approximately 10% of all drug adverse events in the United States, including those that lead to hospitalization (4–5). Novel treatments are needed for all forms of GC excess (6–9).

11β-hydroxysteroid dehydrogenase type 1 (HSD-1) inhibition is a potential new therapeutic approach for conditions of GC excess. HSD-1 converts inactive GC (e.g., cortisone, prednisone) to their active congeners (e.g., cortisol, prednisolone) intracellularly in tissues in which cortisol has physiological roles (e.g., liver, adipose, muscle, skin, bone, brain) and excess cortisol and GC medications are associated with morbidity. HSD-1 is half of a cyclic metabolic pathway.

Active GC (including corticosterone (CORT)) are inactivated in tissues in which they are acutely toxic (e.g., kidney). Inactive GC re-enter circulation where, compared to active GC, they are less protein-bound and thus preferentially enter tissues. HSD-1 is a key tissue-specific regulator of GC levels in human, mouse, and other species, as a substantial portion of active intracellular GC that has access to intracellular receptors is made by HSD-1 from inactive GC (10). Clinical and non-clinical evidence support the potential of HSD-1 inhibition for treatment of GC excess (11–18). Most relevantly to this study, the HSD-1 knockout mouse resisted metabolic, muscular, and dermal morbidity associated with CORT administration for 5 weeks (13). We now show that a HSD-1 inhibitor provides similar benefit as genetic knockout.

SPI-62 (formerly ASP3662) is a potent and selective HSD-1 inhibitor in Phase 2 clinical development for treatment of ACTH-dependent Cushing’s syndrome, ACS, and, in combination with prednisolone, polymyalgia rheumatica. In prior clinical trials (19–21), it was generally well tolerated, demonstrated human pharmacokinetics consistent with once-daily dosing without dependence on timing around meals, and durable human liver and brain HSD-1 inhibition. SPI-62 has also demonstrated human adipose HSD-1 inhibition in a recently completed clinical trial (<u>NCT05409027</u>). In patients with painful diabetic peripheral neuropathy, SPI-62 was associated with decreases, compared to placebo, on glycated hemoglobin, glucose, cholesterol, and triglycerides (22). SPI-62 inhibits mouse HSD-1 with a Ki of 2.6 nM (23).

In this study, CORT was administered in drinking water for 5 weeks to mice, who were randomized to receive three different regimens of SPI-62 or matching vehicle. The SPI-62 regimens were selected to provide >90% HSD-1 inhibition for a limited portion (1 mg/kg QD), most (10 mg/kg QD), or all (10 mg/kg BID) of the day. A control group received neither CORT nor SPI-62. CORT adverse effects on insulin sensitivity, whole-body fat and muscle composition, fat depot and muscle weights, grip strength, dermal thickness, skin structure, and anxiety behavior were planned assessments. Incidental findings on food consumption and weight are also reported.

## Methods

### Animals

Animal use was approved by Institutional Animal Care and Use Committee, Qidong Site (IACUC-QD), WuXi Corporate Committee for Animal Research Ethics (WX-CCARE), WuXi AppTec Co., Ltd. Male C57BL/6 mice with age of 6 to 8 weeks and weight of 19 to 25 g at the start of the study (Lingchang Laboratory Animal Company, Shanghai, China) were housed two per cage and provided food and water *ad libitum*. Temperature and humidity were controlled at 23 ± 2°C and 50 ± 5%. The vivarium was maintained on a 12:12 light:dark cycle with lights on at 7 am.

### Single dose pharmacokinetics

The total number of mice was 54, n = 3 per SPI-62 dose per time point (except that the same animals were used for the 1 and 24 h time points). SPI-62 (Sparrow Pharmaceuticals, Portland, Oregon, USA) was suspended in 0.5% hydroxypropyl methylcellulose (HPMC; Adamas Reagent Company, Shanghai, China) at concentrations of 0.1, 0.3, and 1 mg/mL and administered by oral gavage 10 mL/kg. Whole blood was collected in sodium heparin tubes at 1, 2, 4, 6, 8, 12, and 24 hours after the SPI-62 dose, processed to plasma, mixed 1:1 with 20 mM sodium citrate pH 4.2, and frozen within 2 hours of collection.

A sample volume of 15.0 μL of diluted plasma was aliquoted into a 1.2 mL 96-well plate. 15 μL of internal standard solution (SPI-62-D9 100 ng/mL in water:acetonitrile, 50:50), 50 μL of water:formic acid, 100:0.2, and 600 μL of methyl tert-butyl ether were added to each well. The plates were capped, vigorously shaken, vortex-mixed, and centrifuged. A 300 μL aliquot of the organic phase was transferred to a new plate, evaporated to dryness under nitrogen at ∼ 40°C, and reconstituted in 240 μL of water:methanol:formic acid (50:50:0.1).

Samples were analyzed on a Waters Acquity liquid chromatograph interfaced with a Thermo Scientific TSQ Vantage triple quadrupole mass spectrometer with ESI ionization. Each extracted sample was injected (10 μL) onto a Waters Acquity BEH Phenyl column (2.1 x 50 mm; 1.7 μm) equilibrated at 45°C. Mobile Phase A was water:formic acid, 100:0.1. Mobile Phase B was methanol:formic acid, 100:0.1. The LC flow rate was 0.400 mL/min. The LC gradient was as follows: 50.0%:50.0% A:B from 0.00 to 0.50 minutes, 35.0%:65.0% A:B from 1.50 to 3.00 minutes, and 50.0%:50.0% A:B from 3.25 to 4.25 minutes. The expected retention time for SPI-62 was 1.90 minutes. The precursor and product *m/z* were 425.099 and 262.123. The assay showed ≤ 2.5% intra-run bias for SPI-62 calibration standards 10-10,000 ng/mL and ≤ 7.7% intra-run bias for SPI-62 quality control samples 10-7,500 ng/mL.

### Regimen selection

Contemporaneous plasma SPI-62 concentrations and prednisolone:prednisone ratios in plasma and brain were available to us from an unpublished prior study in which SPI-62 and prednisone were administered to mouse (data on file). We converted the prednisolone:prednisone ratios to percent HSD-1 inhibition, with the value observed in animals who received vehicle for SPI-62 set as 0%. Percent HSD-1 inhibition was modeled on SPI-62 concentrations using the Fit Michaelis-Menten option of the Nonlinear Modeling platform in JMP 17.1.0 software (SAS Institute, Cary NC). IC50, IC80, and IC90 were estimated by inverse prediction.

Plasma concentration-time profiles from the single dose pharmacokinetics study were fit utilizing the Fit Biexponential option of the Fit Curve platform in JMP 17.1.0 software (SAS Institute, Cary NC), the option that provided the best fit for the observed data. The approximate duration that SPI-62 regimens would maintain plasma levels above IC50, IC80, or IC90 were estimated visually.

The CORT regimen was based on prior demonstration of association with multiple adverse events of interest (13).

### Mouse model of glucocorticoid excess

The total number of mice was 70, n = 14 per group. Dosing (Table 1) was on Days 1 through 35 at 10 am (Groups 1 through 4) or at 10 am and 5 pm (Group 5). CORT (Bide Pharmatech, Shanghai, China) was dissolved in 0.66% of final volume ethanol (Sigma-Aldrich Trading Company, Shanghai, China) then further dissolved in drinking water of the mice at a final concentration of 100 μg/mL. Drinking water contained 0.66% v/v ethanol. CORT and drinking water were prepared daily. SPI-62 (Sparrow Pharmaceuticals, Portland, Oregon, USA) was suspended in 0.5% HPMC (Adamas Reagent Company, Shanghai, China) at concentrations of 0.1 and 1 mg/mL and administered by oral gavage 10 mL/kg. SPI-62 and 0.5% HPMC were prepared fresh for each dose.

**Table 1.** Dosing schedules for the experimental groups (n = 14 per group)

| **Group** | **Test**<br>**Article 1** | **Route** | **Dosing Schedule** | **Test**<br>**Article 2** | **Route** | **Dosing Schedule** |
| --- | --- | --- | --- | --- | --- | --- |
| 1 | Vehicle | p.o. | 0.5% HPMC<br>(Day 1 thru Day 35) | Water | In drinking<br>water | 0.66% ethanol<br>(Day 1 thru Day 35) |
| 2 | Vehicle | p.o. | 0.5% HPMC<br>(Day 1 thru Day 35) | CORT | In drinking<br>water | 100 µg/mL<br>(Day 1 thru Day 35) |
| 3 | SPI-62 | p.o. | 1 mg/kg/day<br>(Day 1 thru Day 35, QD) | CORT | In drinking<br>water | 100 µg/mL<br>(Day 1 thru Day 35) |
| 4 | SPI-62 | p.o. | 10 mg/kg/day<br>(Day 1 thru Day 35, QD) | CORT | In drinking<br>water | 100 µg/mL<br>(Day 1 thru Day 35) |
| 5 | SPI-62 | p.o. | 10 mg/kg/day<br>(Day 1 thru Day 35, BID) | CORT | In drinking<br>water | 100 µg/mL<br>(Day 1 thru Day 35) |

### In-life procedures (Figure 1)

A viability check for mortality/moribundity was performed twice daily. Detailed clinical observations, including ophthalmology, were performed weekly. Body weight was measured daily to enable weight-based SPI-62 administration. Food consumption was measured twice weekly for pair-housed animals and reported weekly as a combination of the data from the two measurements during that week.

On Days 1 (Baseline), 15, 29, and 35, mice were food restricted (fasted) for 5 hours from 7 am until noon. Blood samples were collected at noon by nicking the tail vein. Whole blood was collected in heparin sodium (5 mg/mL, 2 μL for 50 μL blood) tubes and centrifuged to plasma within 2 hours of collection. Glucose was measured using a glucometer (OneTouch UltraVue, LifeScan, Milpitas, California, USA) using about 10 μL of blood. Insulin was measured in the remaining blood (40-50 μL) by enzyme-linked immunosorbent assay (MilliporeSigma, Burlington, Massachusetts, USA, catalog #EZRMI-13K). Homeostatic model assessment of insulin resistance (HOMA-IR) was calculated according to the formula:

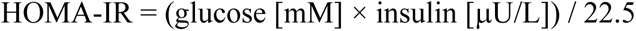

**Figure 1.**
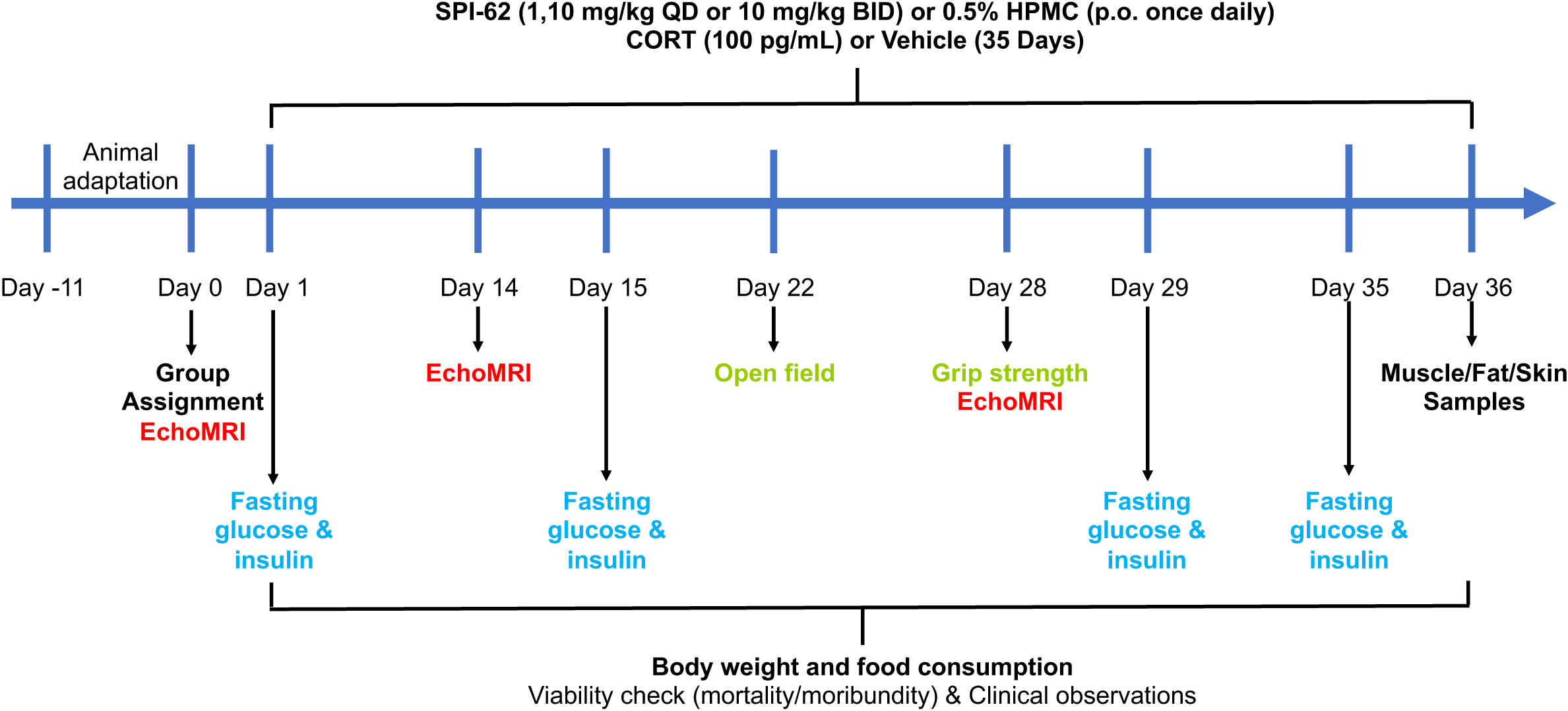
Study schematic (n = 14 per group)

An EchoMRI-130H™ body composition analyzer (Mediso, Budapest, Hungary) was used to measure whole-body fat and muscle tissue according to instructions provided by the manufacturer on Days 0 (baseline), 14, and 28.

Grip strength was measured using a digital grip strength meter (BIOSEB, Vitrolles, France, model BIO-GS3) on Day 28. The mouse was lowered over the grid with the torso kept horizontal and allowing only its forepaws to grasp the grid before measurements were taken. Then, the mouse was pulled gently by its tail while it gripped the grid and the torso remained horizontal. The maximal grip strength value was recorded. This procedure was repeated a total of 3 times per mouse.

Ambulation behavior was assessed in an open field maze on Day 22. A 27 × 27 × 20.3 cm Med Associates box chamber (Lanyuao Industry and Trade Company, Shanghai, China) with 16 infrared and photoemitters on each side of the box and two rows of emitters and detectors on two sides to enable assessment of rearing. Mice were placed in the center of the test chamber and allowed to move freely for 10 minutes. For result analysis, the chamber was divided into central (9 × 9 cm) and peripheral zones. Times spent in each zone, distance traveled, and number of rearings were recorded and analyzed using ANYmaze60000 software (Stoelting, Wood Dale, Illinois, USA, version 4.99z).

### Post-mortem procedures (Figure 1)

Gonadal, subcutaneous, retroperitoneal, and mesenteric fat, quadriceps, and tibialis anterior were excised and weighed.

Skin from mouse back was fixed in 4% paraformaldehyde. Samples were subsequently paraffin embedded. 8-μm sections were prepared on a HISCORE BIOCUT microtome (Leica Microsystems, Wetzlar, Germany). Sections were stained with hematoxylin and eosin. Skin thickness (dermal layer) was quantified using ImageJ software (National Institutes of Health, Bethesda, Maryland, USA, version 1.52p). Representative images were used for qualitative assessment of dermal structure.

### Statistical methods

Animals were assigned randomly to achieve matched mean body weight in each experimental group. Data were collected and processed randomly. Investigators responsible for dosing and data collection were blinded to the randomization sequence. No data were excluded. Absence of non-normal data distributions and outliers was assessed visually.

Formal power calculations were not performed. A study of CORT effects in HSD-1 knockout mouse (13) indicated that, under hypotheses of effect sizes associated with SPI-62 equal to those observed with HSD-1 knockout, statistically significant differences should be achieved with the selected sample size on endpoints of insulin sensitivity, body composition, grip strength, fat and muscle weight, and dermal thickness. That was not so for food consumption, which was not reported in that prior study (13) and had only 7 data per group per time point in our study as it was measured for pairs of co-housed animals. Our study was not designed for statistical testing on food consumption and so none was conducted. No numerical or statistical effect of CORT on body weight was observed in that prior study. With an anticipated effect size of zero statistical testing for a difference between groups was not justifiable. Dermal structure was not quantified. Statistical analyses were conducted using Prism (GraphPad Software, San Diego, California, USA, version 7.0.0.159). Glucose, insulin, HOMA-IR, whole-body fat and muscle content were analyzed by analysis of covariance (ANCOVA) with baseline value as the covariate. Grip strength, open field test parameters, and post-mortem fat depot and muscle weights, for which baseline data were not available, were analyzed by analysis of variance (ANOVA). A two-sided alpha level of 0.05 was used for each analysis without regard for multiple testing.

## Results

### Concentration-response modeling of HSD-1 inhibition

Each of plasma (whole body) and brain HSD-1 inhibition were modeled as dependent variables with plasma SPI-62 as the independent variable (Figure 2). The confidence interval of mean maximum HSD-1 inhibition (Imax) included 100% for both plasma and brain. The Michaelis constant (Km) and 50, 80, and 90% inhibitory constants (IC50, IC80, IC90) were somewhat higher for brain than for plasma (Table 2).

**Figure 2.**
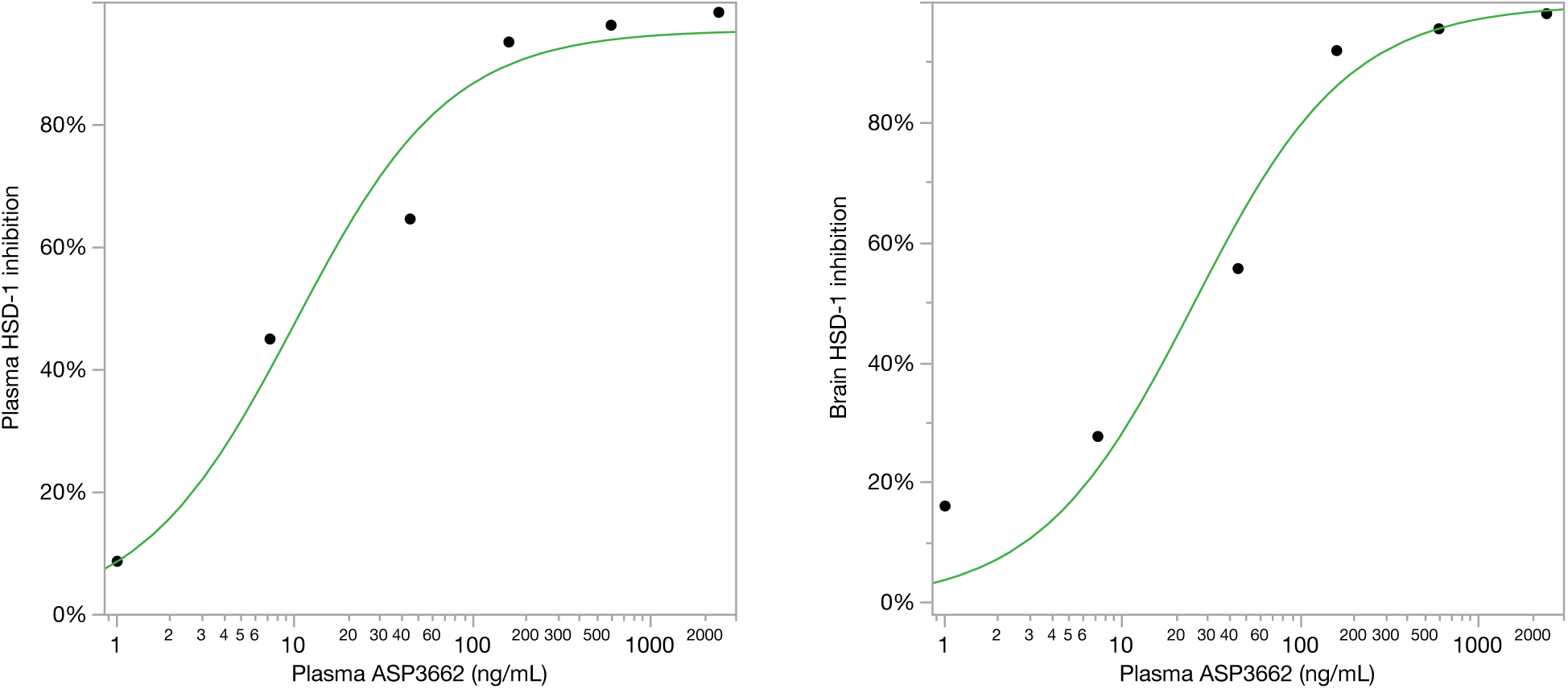
Michaelis-Menten relationships of plasma and brain HSD-1 inhibition to SPI-62 plasma concentrations in mouse. Plasma SPI-62 concentrations and prednisolone:prednisone ratios in plasma and brain from a prior study, in which SPI-62 and prednisone were administered to mouse, were used. Prednisolone:prednisone ratios were converted to percent HSD-1inhibition, with the value observed in animals who received vehicle for SPI-62 set as 0%.

**Table 2.**
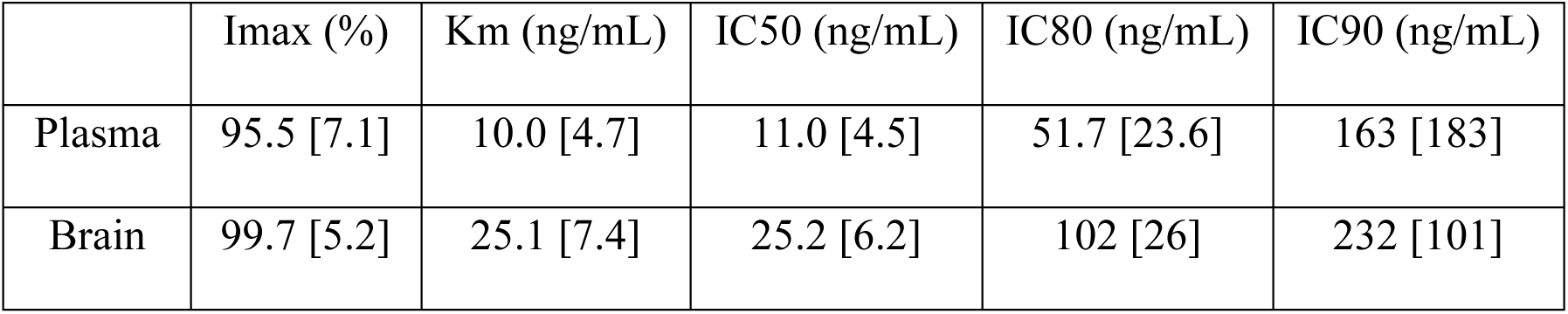
Mean [SE] Michaelis-Menten model parameters of plasma and brain HSD-1 inhibition in mouse.

|  | Imax (%) | Km (ng/mL) | IC50 (ng/mL) | IC80 (ng/mL) | IC90 (ng/mL) |
| --- | --- | --- | --- | --- | --- |
| Plasma | 95.5 [7.1] | 10.0 [4.7] | 11.0 [4.5] | 51.7 [23.6] | 163 [183] |
| Brain | 99.7 [5.2] | 25.1 [7.4] | 25.2 [6.2] | 102 [26] | 232 [101] |

### Single dose pharmacokinetics and regimen selection

SPI-62 plasma concentration-time profiles were best fit by a biexponential curve among several models. Guidelines corresponding to brain IC50, IC80, and IC90 were superimposed on the profiles to enable approximation of the duration that various regimens would maintain HSD-1 inhibition thresholds (Figure 3). Only brain inhibitory constants were used because they were somewhat higher than those for plasma, and the planned study included an endpoint (ambulatory behavior in an open field maze) that would rely on brain HSD-1 inhibition. Our intention was to select a low, medium, and high doses that would maintain IC90 for a limited portion, most, and all, of a day. Approximate durations that selected SPI-62 regimens were expected to maintain HSD-1 inhibition thresholds are presented (Table 3).

**Figure 3.**
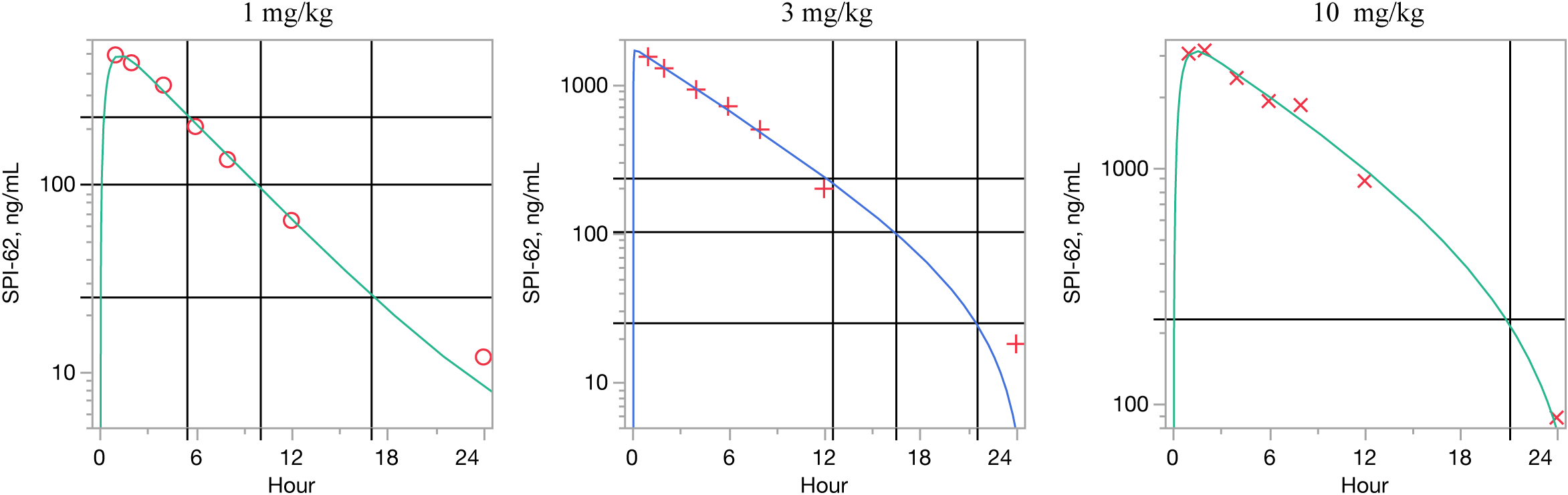
SPi-62 plasma concentration-time curves in mouse. Horizontal lines indicate IC50, IC80, and IC90 for HSD-1 inhibition in brain.

**Table 3.** Approximate duration that SPI-62 regimens selected for mouse Cushing’s syndrome model were expected to maintain HSD-1 inhibition thresholds.

| Regimen | Duration above threshold (h / 24 h) |  |  |
| --- | --- | --- | --- |
|  | IC90 | IC80 | IC50 |
| 1 mg/kg QD | 5.5 | 10 | 17 |
| 10 mg/kg QD | 21 | 24 |  |
| 10 mg/kg BID | 24 |  |  |

### Insulin sensitivity

The mean [SD] baseline HOMA-IR of animals was 0.60 [0.23] mmol glucose/μU insulin and did not differ between groups. CORT was associated with markedly increased HOMA-IR, compared to controls, at each of Day 15, 29, and 35 (Table 4). HOMA-IR was numerically similar between Vehicle + Water and 10 mg/kg BID + CORT groups on Days 15 and 29. Intermediate values were observed for the 1 mg/kg QD + CORT and 10 mg/kg QD + CORT groups.

**Table 4.** Insulin resistance in mice administered CORT and SPI-62. Mice (n = 14 per group) received corticosterone in drinking water and SPI-62 or vehicle by gavage for 35 days. Data were analyzed by ANCOVA with baseline as the covariate.

| Group | HOMA-IR (mmol glucose/μU insulin) | | | | | |
| --- | --- | --- | --- | --- | --- | --- |
|  | Day 15 |  | Day 29 |  | Day 35 |  |
|  | LSM | SE | LSM | SE | LSM | SE |
| Vehicle + Water | 0.78 | 0.46 | 0.73 | 0.37 | 1.02 | 3.86 |
| Vehicle + CORT | 4.02<sup>a</sup> | 0.47 | 5.71<sup>a</sup> | 0.38 | 23.71<sup>a</sup> | 4.01 |
| 1 mg/kg QD + CORT | 3.73<sup>a</sup> | 0.44 | 3.56<sup>ab</sup> | 0.35 | 8.16<sup>b</sup> | 3.73 |
| 10 mg/kg QD + CORT | 1.41<sup>b</sup> | 0.44 | 1.88<sup>ab</sup> | 0.35 | 2.28<sup>b</sup> | 3.72 |
| 10 mg/kg BID + CORT | 0.94<sup>b</sup> | 0.44 | 0.94<sup>b</sup> | 0.36 | 2.16<sup>b</sup> | 3.74 |

<sup>a</sup>P < 0.05 v Vehicle + Water; <sup>b</sup>P < 0.05 v Vehicle + CORT

### Body composition

The mean [SD] baseline whole-body muscle content of animals was 0.87 [0.05] (on a 0-1 scale) and did not differ between groups. CORT was associated with markedly decreased muscle content, compared to controls, on Days 14 and 28 (Table 5). Estimated muscle content was numerically similar between Vehicle + Water for 10 mg/kg QD + CORT on Day 14 and 10 mg/kg BID + CORT on each of Day 14 and 28. Intermediate values were observed for the 1 mg/kg QD + CORT and 10 mg/kg QD + CORT groups.

**Table 5.**
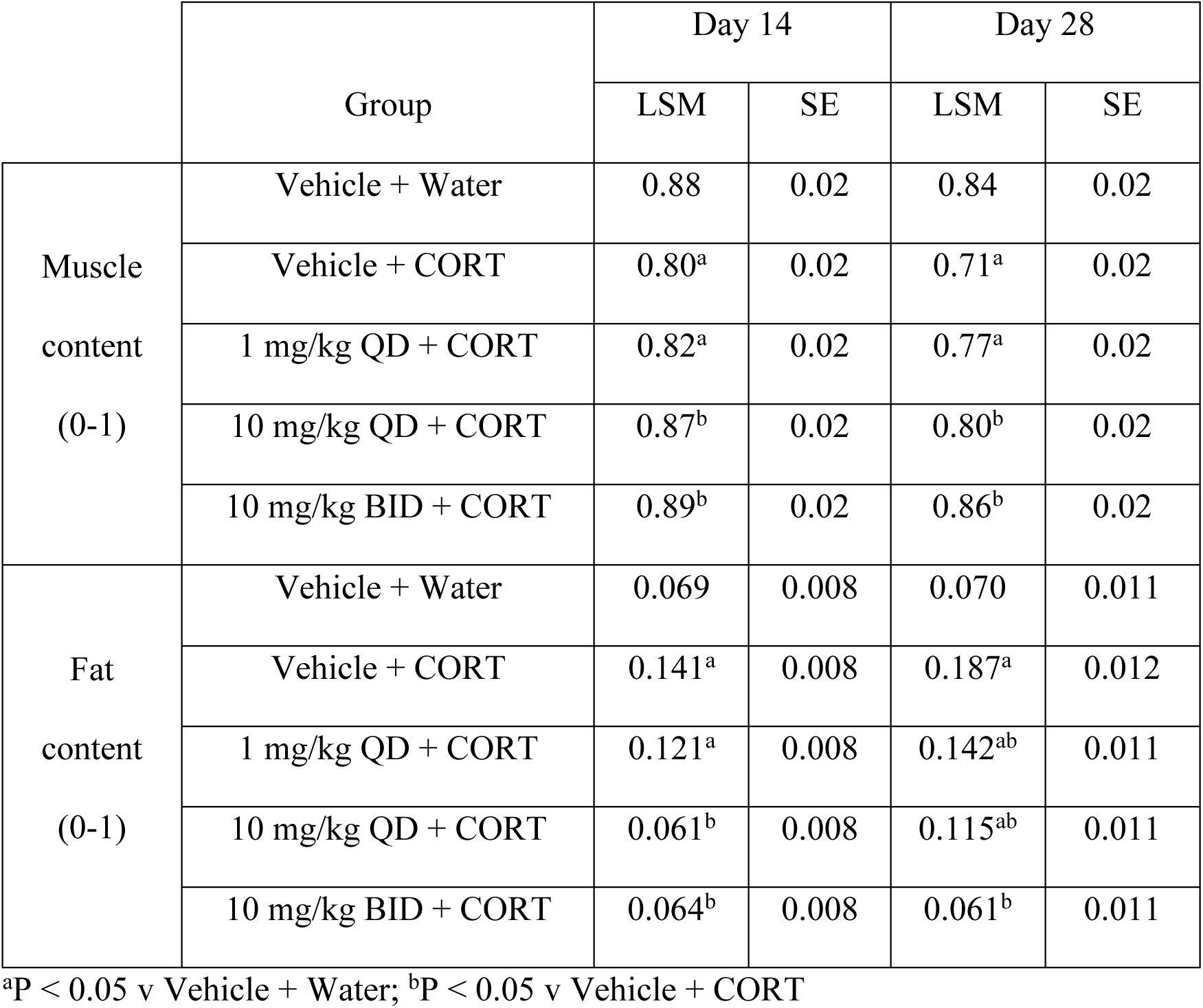
Whole-body muscle and fat content in mice administered CORT and SPI-62. Mice (n = 14 per group) received corticosterone in drinking water and SPI-62 or vehicle by gavage for 35 days. Data were analyzed by ANCOVA with baseline as the covariate.

The mean [SD] baseline whole-body fat content of animals was 0.074 [0.032] and did not differ between groups. CORT was associated with markedly increased fat content, compared to controls, at each of Day 14 and 28 (Table 5). Estimated fat content was numerically similar between Vehicle + Water for 10 mg/kg QD + CORT on Day 14 and 10 mg/kg BID + CORT on each of Day 14 and 28. Intermediate values were observed for the 1 mg/kg QD + CORT and 10 mg/kg QD + CORT groups.

### Grip strength

Mice administered CORT showed numerically decreased forelimb grip strength on Day 28. The Vehicle + CORT group mean [SD] was 3.77 [0.41] compared to 4.29 [0.44] g /g body weight for the Vehicle + Water group; the difference was not statistically significant. Grip strength in groups who received both drugs were 4.98 [1.09], 4.94 [0.72], and 5.40 [0.70] g /g body weight for 1 mg/kg QD + CORT, 10 mg/kg QD + CORT, and 10 mg/kg BID + CORT groups (each P < 0.05 v Vehicle + Water and P < 0.05 v Vehicle + CORT).

### Fat and muscle weights

Post-mortem fat and muscle weights were normalized by body weight.

CORT was associated with substantial increases on Day 36 in the weight of gonadal, subcutaneous, retroperitoneal, and mesenteric fat depots (Table 6). The gonadal, subcutaneous, and retroperitoneal fat depot weights showed similar numerical values between Vehicle + Water and 10 mg/kg BID + CORT groups, but the mesenteric fat depot did not. Intermediate values were observed for the 1 mg/kg QD + CORT and 10 mg/kg QD + CORT groups.

**Table 6.**
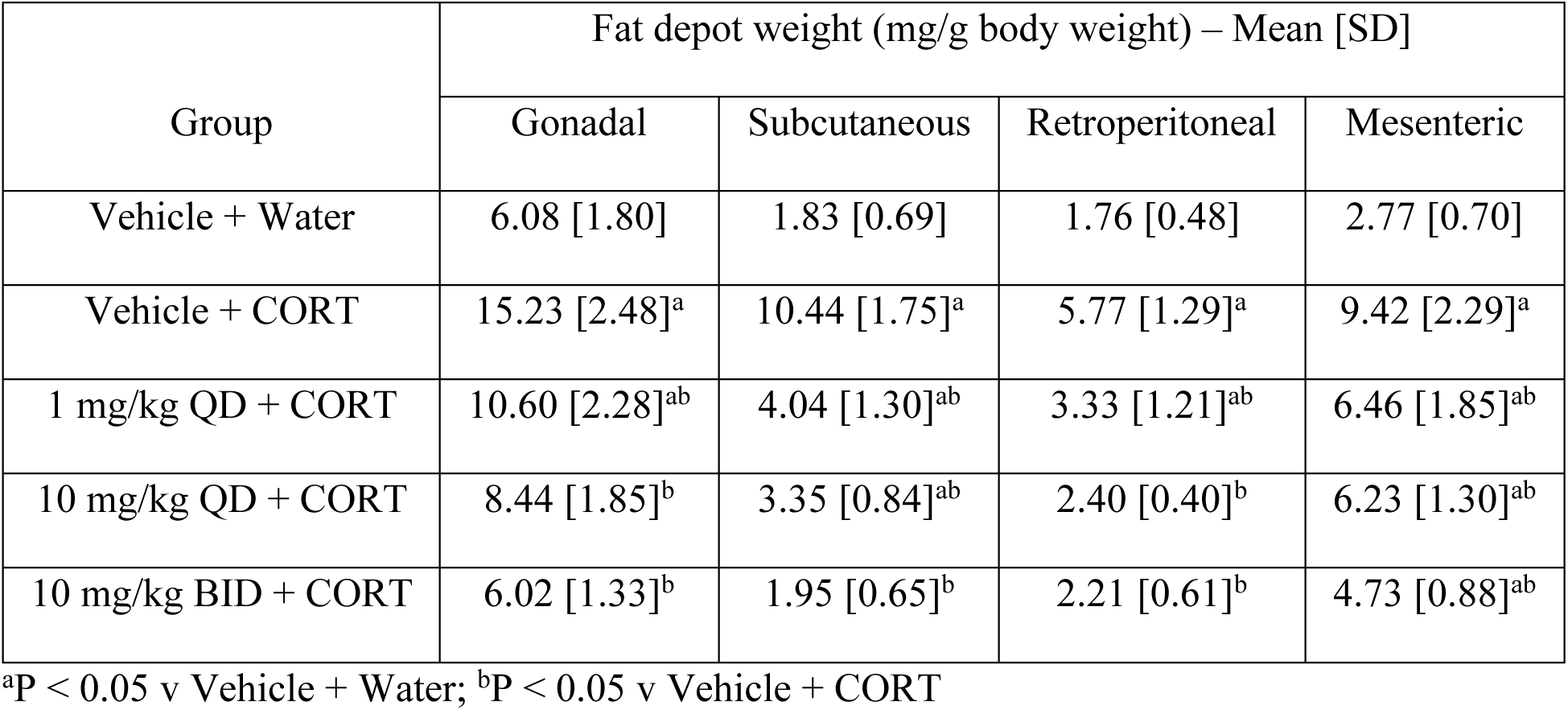
Post-mortem (Day 36) fat depot weights in mice administered CORT and SPI-62. Mice (n = 14 per group) received corticosterone in drinking water and SPI-62 or vehicle by gavage for 35 days. Data were analyzed by ANOVA.

| Group | Fat depot weight (mg/g body weight) – Mean [SD] |  |  |  |
| --- | --- | --- | --- | --- |
|  | Gonadal | Subcutaneous | Retroperitoneal | Mesenteric |
| Vehicle + Water | 6.08 [1.80] | 1.83 [0.69] | 1.76 [0.48] | 2.77 [0.70] |
| Vehicle + CORT | 15.23 [2.48]<sup>a</sup> | 10.44 [1.75]<sup>a</sup> | 5.77 [1.29]<sup>a</sup> | 9.42 [2.29]<sup>a</sup> |
| 1 mg/kg QD + CORT | 10.60 [2.28]<sup>ab</sup> | 4.04 [1.30]<sup>ab</sup> | 3.33 [1.21]<sup>ab</sup> | 6.46 [1.85]<sup>ab</sup> |
| 10 mg/kg QD + CORT | 8.44 [1.85]<sup>b</sup> | 3.35 [0.84]<sup>ab</sup> | 2.40 [0.40]<sup>b</sup> | 6.23 [1.30]<sup>ab</sup> |
| 10 mg/kg BID + CORT | 6.02 [1.33]<sup>b</sup> | 1.95 [0.65]<sup>b</sup> | 2.21 [0.61]<sup>b</sup> | 4.73 [0.88]<sup>ab</sup> |

<sup>a</sup>P < 0.05 v Vehicle + Water; <sup>b</sup>P < 0.05 v Vehicle + CORT

CORT was associated with substantial decreases on Day 36 in the weight of quadriceps and tibialis anterior (Table 7). The means of Vehicle + Water, 10 mg/kg QD + CORT, and 10 mg/kg BID + CORT were numerically similar for tibialis anterior but not quadriceps. Intermediate values were observed for the 1 mg/kg QD + CORT and 10 mg/kg QD + CORT groups.

**Table 7.** Post-mortem (Day 36) muscle weights in mice administered CORT and SPI-62. Mice (n = 14 per group) received corticosterone in drinking water and SPI-62 or vehicle by gavage for 35 days. Data were analyzed by ANOVA.

| Group | Muscle weight (mg/g body weight) – Mean [SD] |  |
| --- | --- | --- |
|  | Quadriceps | Tibialis anterior |
| Vehicle + Water | 6.99 [1.14] | 2.25 [0.24] |
| Vehicle + CORT | 3.19 [0.86]<sup>a</sup> | 1.47 [0.30]<sup>a</sup> |
| 1 mg/kg QD + CORT | 3.98 [0.77]<sup>ab</sup> | 1.60 [0.18]<sup>a</sup> |
| 10 mg/kg QD + CORT | 5.33 [[0.70]<sup>ab</sup> | 2.14 [0.32]<sup>b</sup> |
| 10 mg/kg BID + CORT | 5.90 [0.83]<sup>ab</sup> | 2.36 [0.32]<sup>b</sup> |

<sup>a</sup>P < 0.05 v Vehicle + Water; <sup>b</sup>P < 0.05 v Vehicle + CORT

### Dermal thickness and structure

CORT was associated with a substantial decrease on Day 36 in dermal thickness. The mean [SD] of mice administered Vehicle + CORT was 242.8 [58.2] μm compared to 339.1 [88.2] μm in Vehicle + Water mice (P < 0.05). Mice administered SPI-62 + CORT showed numerical improvement on this parameter: 274.0 [59.2] μm for 1 mg/kg QD + CORT, 268.0 [78.6] μm for 10 mg/kg QD + CORT, and 275.9 [90.2] μm for 10 mg/kg BID + CORT. The 1 mg/kg QD and 10 mg/kg BID group means were statistically significant (P < 0.05) compared to that of Vehicle + Water.

CORT administration demonstrated compaction of the dermal layer consistent with lowering of dermal collagen content. SPI-62 reversed the adverse effect of CORT in a dose-dependent manner, such that the skin structure of mice administered the highest dose was visually similar as that of controls (Figure 4).

**Figure 4.**
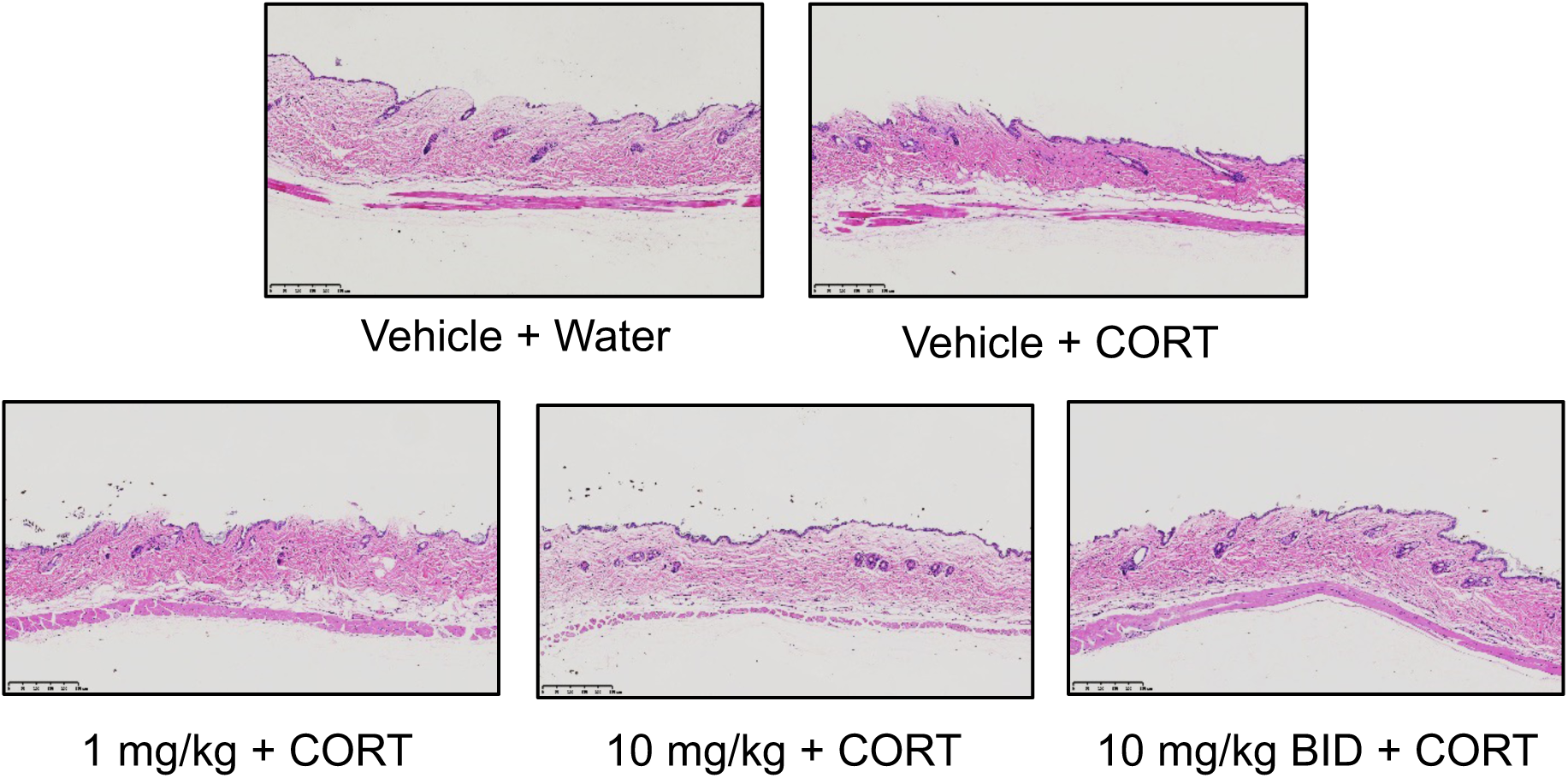
Hematoxylin-and eosin-stained dermal cross-sections in mice administered CORT and SPI-62. Mice (n = 14 per group) received corticosterone in drinking water and SPI-62 or vehicle by gavage for 35 days. Skin from the back was fixed and stained post-mortem on Day 36.

### Open field test

No differences were observed between the vehicle + water and vehicle + CORT groups for time spent in the central zone (28.57 [10.55] v 32.11 [9.68] sec), distance traveled in the central zone (2.69 [1.05] v 2.33 [0.81] m), total distance traveled (31.80 [6.44] v 28.4 [4.46] m), or number of rearings (53 [15] v 47 [10]). As there was no observed CORT effect to modulate, results for groups that received SPI-62 are not shown.

### Clinical observations

Two (of 14) animals in the Vehicle + CORT group died, one each on Day 11 and Day 19. The causes of death were not determined. One (of 14) animal in the Vehicle + Water group died due to accidental gavage injury on Day 29. All 42 animals that received SPI-62 survived until sacrifice on Day 36.

CORT was associated with numerically increased food consumption throughout the study. Mean food consumption in the groups who received both drugs was numerically more like the Vehicle + Water group than to the Vehicle + CORT group (Table 8).

**Table 8.** Observed group mean [standard deviation] food consumption in mice administered CORT and SPI-62. Mice (n = 7 co-housed pairs per group received corticosterone in drinking water and SPI-62 or vehicle by gavage for 35 days. Hypothesis testing was neither pre-specified nor conducted (see Methods).

|  | Food consumption (g/mouse/day) |  |  |  |  |  |  |  |  |  |
| --- | --- | --- | --- | --- | --- | --- | --- | --- | --- | --- |
|  | Vehicle +<br>Water |  | Vehicle +<br>CORT |  | 1 mpk QD +<br>CORT |  | 10 mpk QD +<br>CORT |  | 10 mpk BID +<br>CORT |  |
| Day | Mean | SD | Mean | SD | Mean | SD | Mean | SD | Mean | SD |
| 4 | 2.99 | 0.42 | 3.75 | 0.35 | 3.26 | 0.24 | 3.37 | 0.42 | 3.36 | 0.32 |
| 8 | 3.05 | 0.21 | 3.43 | 0.53 | 2.92 | 0.20 | 2.98 | 0.27 | 2.88 | 0.21 |
| 11 | 3.12 | 0.27 | 3.30 | 0.37 | 3.09 | 0.25 | 3.11 | 0.32 | 2.92 | 0.25 |
| 15 | 2.92 | 0.34 | 3.28 | 0.36 | 3.04 | 0.33 | 2.91 | 0.20 | 2.74 | 0.22 |
| 18 | 3.13 | 0.26 | 3.19 | 0.34 | 2.91 | 0.30 | 3.08 | 0.18 | 3.36 | 0.43 |
| 22 | 3.06 | 0.28 | 3.25 | 0.35 | 3.23 | 0.37 | 3.15 | 0.33 | 2.89 | 0.28 |
| 25 | 3.24 | 0.19 | 3.61 | 0.22 | 3.74 | 0.36 | 3.50 | 0.19 | 3.13 | 0.30 |
| 29 | 2.92 | 0.23 | 3.32 | 0.21 | 2.90 | 0.18 | 2.99 | 0.34 | 2.79 | 0.29 |
| 32 | 3.10 | 0.20 | 3.26 | 0.32 | 2.77 | 0.37 | 3.00 | 0.62 | 3.27 | 0.60 |
| 36 | 3.00 | 0.29 | 3.28 | 0.30 | 3.13 | 0.17 | 3.24 | 0.12 | 2.83 | 0.30 |

The mean [SD] baseline body weight of animals was 22.06 [1.09] g and did not differ numerically between groups. CORT showed an unexpected biphasic effect on body weight (Figure 5 and Table 9): blockade of the body weight gain was observed during the first two weeks (compare baseline to Day 15) followed by accelerated weight gain compared to controls during the next three weeks (compare Day 15 to Day 35). The mice in both groups had numerically similar body weights on Day 35. SPI-62 did not alter the early effect of CORT through Day 15. In contrast, body weight gain acceleration in groups who received both drugs during the third to fifth weeks of the study appeared to be slower. Body weights on Day 35 in those groups were numerically lower than for either Vehicle + Water or Vehicle + CORT groups. A dose-dependent trend was visually apparent (Figure 5).

**Figure 5.**
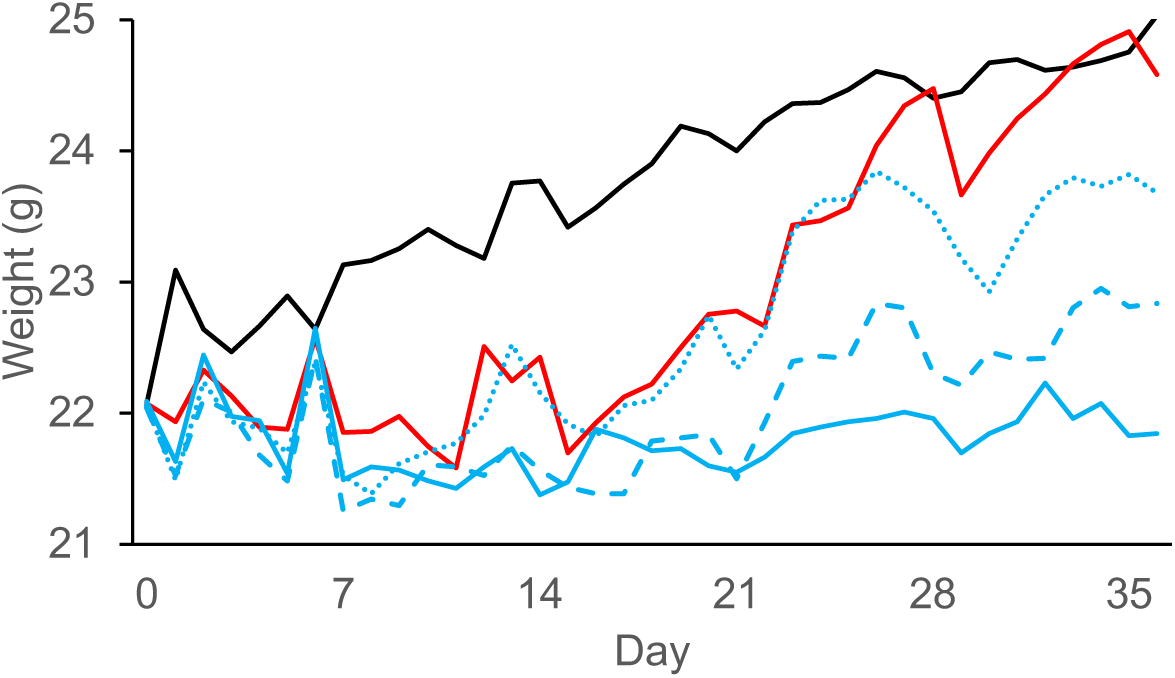
Observed group mean body weights in mice administered CORT and SPI-62. Mice (n = 14 per group) received corticosterone in drinking water and SPI-62 or vehicle by gavage for 35 days. Food consumption was measured twice weekly. Weight was measured daily. No statistical testing was performed. SPI-62 groups are as follows: Vehicle + Water (solid black), Vehicle + CORT (solid red), 1 mg/kg + CORT (dotted blue), 10 mg/kg + CORT (dashed blue), 10 mg/kg BID + CORT (solid blue).

**Table 9.** Observed group means and standard deviations of body weights at Baseline and at Davs 15, 29, and 35 in mice administered CORT and SPI-62. Mice (n = 14 per group received corticosterone in drinking water and SPI-62 or vehicle by gavage for 35 days. Hypothesis testing was neither pre-specified nor conducted.

|  | Body weight (g) |  |  |  |  |  |  |  |
| --- | --- | --- | --- | --- | --- | --- | --- | --- |
| Group | Baseline |  | Day 15 |  | Day 29 |  | Day 35 |  |
|  | Mean | SD | Mean | SD | Mean | SD | Mean | SD |
| Vehicle + Water | 22.1 | 1.12 | 23.4 | 1.71 | 24.4 | 1.72 | 24.8 | 1.69 |
| Vehicle + CORT | 22.1 | 1.06 | 21.7 | 1.88 | 23.7 | 1.91 | 24.9 | 1.80 |
| 1 mpk QD + CORT | 22.1 | 1.04 | 21.9 | 2.00 | 23.2 | 2.80 | 23.8 | 2.96 |
| 10 mpk QD + CORT | 22.0 | 1.09 | 21.4 | 1.10 | 22.2 | 1.32 | 22.8 | 1.21 |
| 10 mpk BID + CORT | 22.1 | 1.29 | 21.5 | 1.29 | 21.7 | 1.51 | 21.8 | 1.50 |

## Discussion

In this study, CORT 100 mg/mL in drinking water administered to mice for 5 weeks resulted in several metabolic (increased insulin resistance, increased adiposity, increased food consumption, accelerated weight gain), muscular (peripheral myoatrophy, weakness), and dermal (thinning, structure change) morbidities that are prominently associated with Cushing’s disease and other forms of GC excess in humans. SPI-62, a potent and selective HSD-1 inhibitor, attenuated all observed CORT adverse effects, generally in a dose-dependent manner.

The CORT-associated adverse effects reported here, except for the biphasic impact on body weight, were previously reported using the identical CORT dose, duration, and mode of administration in the same mouse strain. Further, and consonant with our results, HSD-1 knockout mice did not show those toxicities (13). As regards body weight, the previous authors only reported that CORT-treated and control mice did not differ at the end of their study. We observed the same result on Day 35 between Vehicle + Water and Vehicle + CORT groups.

Because we incidentally had daily weights to enable weight-based SPI-62 dosing, we noticed an unexpected biphasic weight change associated with CORT. Another group has recently observed a similar biphasic weight change in CORT-treated C57BL/6 mice (24). The first phase, in which CORT was associated with decreased body weight gain, appeared not to be affected by SPI-62, and does not translate to humans under conditions of GC excess. We do not have mechanistic hypothesis for decreased weight gain when the animals were hyperphagic and showed insulin resistance. SPI-62 appeared to attenuate the second phase, CORT-associated accelerated weight gain, which is a core concern expressed by patients with endogenous hypercortisolism (25) or who take GC medicines (26). To date, SPI-62 has not been administered for 5 weeks or longer to mice in the absence of CORT, hence, no inference can be made regarding an independent subchronic or chronic effect of SPI-62 on weight in mice.

SPI-62 doses in this study were selected to demonstrate dose-dependent attenuation of CORT adverse effects. HSD-1 inhibitor clinical trial results suggest that to achieve maximum clinical benefit, complete and continuous HSD-1 inhibition is necessary. This concept is best supported in the clinical literature for glycemic control with glycated hemoglobin (HbA1c) as the endpoint (27). The low, medium, and high doses in our study were expected to maintain >90% HSD-1 inhibition for a limited portion, most, and all, of the day. The selected regimens were effective for the intended purpose: clear dose-dependent attenuation was observed on all CORT-associated morbidity observed in the study. Overall, the results suggest that clinical trials of SPI-62 should utilize a dose that achieves full HSD-1 inhibition throughout a dose interval.

On grip strength, mice administered SPI-62 showed superior results compared to the control group. Neither muscle mass nor body muscle content increased. Speculatively, as grip strengths were normalized by body weight, that might reflect that grip strength changes in mice administered SPI-62 were less than proportional to the decreased weight gain observed in those animals.

Except for behavior in the open field maze, CORT was associated with expected morbidity in the mouse model of Cushing’s syndrome. Open field test results can be influenced by subtle variations of temperature, humidity, lighting, noise, odor, and handling (28s) such that even careful experimentation will not end in the same results every time. Those factors could account for lack of observed difference between Vehicle + Water and Vehicle + CORT group. The results of this study support potential that SPI-62, or HSD-1 inhibitors in general, might alleviate certain symptoms of Cushing’s disease and other forms of GC excess in human. Limitations of this study include that several important adverse effects of GC excess(e.g., osteoporosis, neuropsychiatric effects, glaucoma) were assessed.

This study adds to the body of evidence that HSD-1 inhibition can prevent adverse effects of GC excess in general. Clinical trials in patients with ACTH-dependent Cushing’s syndrome, ACS, or who rely on long-term GC therapy are needed to evaluate whether SPI-62 has clinical benefit.

## Data Access

Restrictions apply to the availability of some data generated or analyzed during this study because they were used under license. The corresponding author will on request detail the restrictions and any conditions under which access to some data may be provided.

## Abbreviations

ACS: autonomous cortisol secretion
BID: twice daily
CI: confidence interval
CORT: corticosterone
GC: glucocorticoid(s)
HOMA-IR: homeostatic model assessment of insulin resistance
HPMC: hydroxypropyl methylcellulose
HSD-1: 11β3-hydroxysteroid dehydrogenase type 1
IC50, IC80, IC90: plasma concentrations associated with 50%, 80%, 90% HSD-1 inhibition
Imax: maximum HSD-1 inhibition
Km: Michaelis constant
LSM: least squares mean
QD: once daily
SD: standard deviation
SE: standard error

## References

1. Melmed S. Pituitary-Tumor Endocrinopathies. N Engl J Med. 2020;382(10):937–950. doi:10.1056/NEJMra1810772

2. Limumpornpetch P, Morgan AW, Tiganescu A, et al. The Effect of Endogenous Cushing Syndrome on All-cause and Cause-specific Mortality. J Clin Endocrinol Metab. 2022;107(8):2377–2388. doi:10.1210/clinem/dgac265

3. Ebbehoj A, Li D, Kaur RJ, et al. Epidemiology of adrenal tumours in Olmsted County, Minnesota, USA: a population-based cohort study. Lancet Diabetes Endocrinol. 2020;8(11):894–902. doi:10.1016/S2213-8587(20)30314-4

4. Sarnes E, Crofford L, Watson M, Dennis G, Kan H, Bass D. Incidence and US Costs of Corticosteroid-Associated Adverse Events: A Systematic Literature Review. Clin Ther. 2011;33(10):1413–1432. doi:10.1016/j.clinthera.2011.09.009

5. Weiss AJ, Elixhauser A, Bae J, Encinosa W. Origin of Adverse Drug Events in U.S. Hospitals, 2011. HCUP Statistical Brief #158. 2013. http://www.hcup-us.ahrq.gov/reports/statbriefs/sb158.pdf.

6. Hinojosa-Amaya J, Cuevas-Ramos D, Fleseriu M. Medical Management of Cushing’s Syndrome: Current and Emerging Treatments. Drugs. 2019;79(9):935–956. doi:10.1007/s40265-019-01128-7

7. Bancos I, Taylor AE, Chortis V, et al. Urine steroid metabolomics for the differential diagnosis of adrenal incidentalomas in the EURINE-ACT study: a prospective test validation study. Lancet Diabetes Endocrinol. 2020;8(9):773–781. doi:10.1016/S2213-8587(20)30218-7

8. Rooke TW. The Quest for Cortisone. Michigan State University Press; 2012.

9. Fardet L, Petersen I, Nazareth I. Prevalence of long-term oral glucocorticoid prescriptions in the UK over the past 20 years. Rheumatology. 2011;50(11):1982–1990. doi:10.1093/rheumatology/ker017

10. Tomlinson JW, Walker EA, Bujalska IJ, et al. 11β-Hydroxysteroid Dehydrogenase Type 1: A Tissue-Specific Regulator of Glucocorticoid Response. Endocr Rev. 2004;25(5):831–866. doi:10.1210/er.2003-0031

11. Tomlinson JW, Draper N, Mackie J, et al. Absence of Cushingoid phenotype in a patient with Cushing’s disease due to defective cortisone to cortisol conversion. J Clin Endocrinol Metab. 2002;87(1):57–62. doi:10.1210/jcem.87.1.8189

12. Arai H, Kobayashi N, Nakatsuru Y, et al. A Case of Cortisol Producing Adrenal Adenoma without Phenotype of Cushing’s Syndrome due to Impaired 11β-Hydroxysteroid Dehydrogenase 1 Activity. Endocr J. 2008;55(4):709–715. doi:10.1507/endocrj.k08e-008

13. Morgan SA, McCabe EL, Gathercole LL, et al. 11β-HSD1 is the major regulator of the tissue-specific effects of circulating glucocorticoid excess. Proc Natl Acad Sci U S A. 2014;111(24):E2482–E2491. doi:10.1073/pnas.1323681111

14. Song Z, Gong Y, Liu H, Ren Q, Sun X. Glycyrrhizin could reduce ocular hypertension induced by triamcinolone acetonide in rabbits. Mol Vis. 2011;17:2056–2064. doi:PMID: 21850181; PMCID: PMC3156820

15. Oda S, Ashida K, Uchiyama M, et al. An Open-label Phase I/IIa Clinical Trial of 11β-HSD1 Inhibitor for Cushing’s Syndrome and Autonomous Cortisol Secretion. J Clin Endocrinol Metab. 2021;106(10):E3865–E3880. doi:10.1210/clinem/dgab450

16. Fenton CG, Doig CL, Fareed S, et al. 11β-HSD1 plays a critical role in trabecular bone loss associated with systemic glucocorticoid therapy. Arthritis Research & Therapy. 2019;21(1):188. doi:10.1186/s13075-019-1972-1

17. Fenton CG, Crastin A, Martin CS, et al. 11β-Hydroxysteroid Dehydrogenase Type 1 within Osteoclasts Mediates the Bone Protective Properties of Therapeutic Corticosteroids in Chronic Inflammation. Int J Mol Sci. 2022;23(13):7334. doi:10.3390/ijms23137334

18. Othonos N, Pofi R, Arvaniti A, et al. 11β-HSD1 inhibition in men mitigates prednisolone-induced adverse effects in a proof-of-concept randomised double-blind placebo-controlled trial. Nat Commun. 2023;14:1025. doi: 10.1038/s41467-023-36541-w

19. Bellaire S, Walzer M, Wang T, Krauwinkel W, Yuan N, Marek GJ. Safety, Pharmacokinetics, and Pharmacodynamics of ASP3662, a Novel 11β-Hydroxysteroid Dehydrogenase Type 1 Inhibitor, in Healthy Young and Elderly Subjects. Clinical and Translational Science. 2019;12(3):291–301. doi:10.1111/cts.12618

20. Gallezot J-D, Nabulsi N, Henry S, et al. Imaging the Enzyme 11β-Hydroxysteroid Dehydrogenase Type 1 with PET: Evaluation of the Novel Radiotracer 11C-AS2471907 in Human Brain. J Nucl Med. 2019;60(8):1140–1146. doi:10.2967/jnumed.118.219766

21. Astellas Clinical Trial Results. Available at https://www.trialsummaries.com/Study/StudyDetails?id=14232&tenant=MT_AST_9011 [accessed 13 October 2023]

22. Fleseriu M, Bancos I, Katz D. A double-blind, randomized, placebo-controlled trial of SPI-62 safety and efficacy for the treatment of Cushing s syndrome. Endocrine Abstracts. May 2021 (Abstract AEP546). doi:10.1530/endoabs.73.AEP546

23. Kiso T, Sekizawa T, Uchino H, Tsukamoto M, Kakimoto S. Analgesic effects of ASP3662, a novel 11β-hydroxysteroid dehydrogenase 1 inhibitor, in rat models of neuropathic and dysfunctional pain. British Journal of Pharmacology. 2018;175(19):3784–3796. doi:10.1111/bph.14448

24. Gado M, Heinrich A, Wiedersich D, et al. Activation of b-adrenergic receptor signaling prevents glucocorticoid-induced obesity and adipose tissue dysfunction in male mice. Am J Physiol Endocrinol Metab. 2023;324:E514–E530. doi: 10.1152/ajpendo.00259.2022

25. Gotch PM. Cushing’s syndrome from the patient’s perspective. Endocrinol Metab Clin North Am. 1994;23(3):607–617. doi:10.1016/S0889-8529(18)30087-2

26. Tieu J, Cheah JTI, Black RJ, et al. Improving benefit-harm assessment of glucocorticoid therapy incorporating the patient perspective: The OMERACT glucocorticoid core domain set. Semin Arthritis Rheum. 2021;51(5):1139–1145. doi:10.1016/j.semarthrit.2021.06.010

27. Rosenstock J, Banarer S, Fonseca VA, et al. The 11β-Hydroxysteroid Dehydrogenase Type 1 Inhibitor INCB13739 Improves Hyperglycemia in Patients With Type 2 Diabetes Inadequately Controlled by Metformin Monotherapy. Diabetes Care. 2010;33(7):1516–1522. doi:10.2337/dc09-2315

28. Tatem KS, Quinn JL, Phadke A, Yu Q, Gordish-Dressman H, Nagaraju K. Behavioral and locomotor measurements using an open field activity monitoring system for skeletal muscle diseases. J Vis Exp. 2014;(91):51785. doi:10.3791/51785

